# Diffusion time-related structure-function coupling reveals differential association with inter-individual variations in body mass index

**DOI:** 10.1101/2023.07.18.549603

**Authors:** Jong Young Namgung, Yeong Jun Park, Yunseo Park, Chae Yeon Kim, Bo-yong Park

**Author notes:** **Corresponding Author:** Bo-yong Park, PhD, Department of Data Science, Inha University, Incheon, Republic of Korea.

## Abstract

Body mass index (BMI) is an indicator of obesity, and recent neuroimaging studies have demonstrated inter-individual variations in BMI to be associated with altered brain structure and function. However, how the structure-function correspondence is altered according to BMI is under-investigated. In this study, we combined structural and functional connectivity using Riemannian optimization with varying diffusion time parameters and assessed their association with BMI. First, we simulated functional connectivity from structural connectivity and generated low-dimensional principal gradients of the simulated functional connectivity across diffusion times, where low and high diffusion times indirectly reflected mono- and polysynaptic communication. We found the most apparent cortical hierarchical organization differentiating between low-level sensory and higher-order transmodal regions in the middle of the diffusion time, indicating that the hierarchical organization of the brain may reflect the intermediate mechanisms of mono- and polysynaptic communications. Associations between the simulated gradients and BMI revealed the strongest relationship when the hierarchical structure was most evident. Moreover, the functional gradient-BMI association map showed significant correlations with the cytoarchitectonic measures of the microstructural gradient and moment features, indicating that BMI-related functional connectome alterations were remarkable in higher-order cognitive control-related brain regions. Finally, transcriptomic association analysis provided potential biological underpinnings, specifying gene enrichment in the striatum, hypothalamus, and cortical cells. Our findings provide evidence that structure-function correspondence is strongly coupled with BMI when hierarchical organization is most apparent, and the associations are related to the multiscale properties of the brain, leading to an advanced understanding of the neural mechanisms related to BMI.

## INTRODUCTION

Obesity is a prevalent condition worldwide, which is easily measured using the body mass index (BMI)[1,2]. Managing body weight is important because a high BMI can lead to health-related problems such as type 2 diabetes, cardiovascular disease, sleep apnea, and comorbidities[3–6]. Moreover, previous studies have found inter-individual variations in BMI to be associated with cognitive function and cell-type-specific metabolic activity[7–10]. However, studies linking BMI to large-scale structural and functional brain networks and neuronal mechanisms are relatively scarce.

Neuroimaging studies based on magnetic resonance imaging (MRI) have revealed differences in the brain morphology and inter-regional brain connectivity related to variations in the BMI. For example, previous studies have found structural alterations in the gray matter and white matter in individuals with a high BMI[11–15] and dysfunction in functional brain networks, and their relationship to abnormal appetite and energy regulation[16–18]. Recent studies have adopted a connectivity analysis to assess interregional functional connectivity based on a correlation analysis, and structural connectivity based on diffusion tractography[19,20]. This graph-theoretical connectivity analysis has been widely adopted to assess the association between the BMI and brain networks[16,17]. However, how structural and functional connectome organizations are simultaneously related to the BMI is relatively under-investigated, although it is evident that brain structure and function are closely intertwined[21–26]. In this study, we aimed to explore the structure-function coupling of the brain and its association with inter-individual variations in the BMI.

Structure-function correspondences have been widely investigated in previous studies by predicting functional connectivity from structural connectivity via biophysical modeling and graph-based network communication models[27–34]. These models are based on synaptic communication, which considers polysynaptic pathways in functional interactions[33–37]. A recent study proposed a Riemannian optimization framework to assess structure-function coupling based on a synaptic communication model[38]. The key idea of this approach is a dimensionality-reduction technique that aims to identify a transformation matrix, which rotates the low-dimensional eigenvectors of structural connectivity to reconstruct functional connectivity. It is governed by a diffusion time parameter that reflects the implications of the polysynaptic pathways[38]. In a previous study, the sensory/motor regions were well predicted at lower diffusion times (i.e., monosynaptic), whereas the higher-order default-mode regions required higher diffusion times (i.e., polysynaptic). Here, we assessed structure-function coupling using the Riemannian optimization framework and associated it with BMI with varying diffusion times. We hypothesized that structurally governed functional brain organization may exhibit differential polysynaptic mechanisms related to obesity-related traits.

Multiscale analyses using cytoarchitectural and transcriptomic data can complement the imaging-based findings. Previous studies integrated large-scale structural or functional brain networks with microcircuit functions and gene expression data[39–44]. For example, our previous work suggested a consolidated framework linking structural connectome alterations in individuals with autism spectrum disorder and neuronal excitation/inhibition imbalance, as well as developmental enrichment of gene expression[45]. Such analyses have been widely adopted to investigate the multiscale properties of shared effects of multiple psychiatric conditions[41,46]. Taken together, this multiscale framework may provide insights into the underlying biological processes related to network-level brain alterations.

In this study, we investigated structure-function coupling using a Riemannian optimization framework [38] and assessed its relationship with inter-individual variations in BMI. Furthermore, we performed multiscale analysis by linking macroscale data to cytoarchitectural and gene expression data. Our work provides insights into the neurobiological aspects of BMI-related structure-function coupling.

## RESULTS

### Structure-function coupling using Riemannian optimization

We opted for a Riemannian optimization framework to predict functional connectivity from structural connectivity [38] (**Fig. 1a**), using multimodal MRI data obtained from the Human Connectome Project (HCP) database[47]. In brief, this technique determines the optimal transformation of structural connectivity to reconstruct functional connectivity in a low-dimensional manifold space by varying the diffusion time (*t*) [38] (see *Methods*). At the global level, the prediction performance was improved as the diffusion time increased, and the higher spatial granularity showed less variability than lower granularities (mean ± standard deviation (SD) correlation coefficients = 0.775 ± 0.024 / 0.810 ± 0.015 / 0.805 ± 0.022 for 200, 300, and 400 parcels at *t* = 1; and 0.870 ± 0.016 / 0.900 ± 0.017 / 0.871 ± 0.022 at *t* = 10; **Fig. 1b**). We repeated the analysis 30 times using different training and test datasets to confirm the stability of the predictive model (mean and 95% confidence interval [CI] of the correlation coefficients = 0.805 (0.8028, 0.8079) / 0.904 (0.9009, 0.9073) at *t* = 1 and 10 for 300 parcels; **Fig. 1b**). We quantified the prediction performance for each brain region; similarly, a greater predictive performance was observed at higher diffusion times (**Fig. 1c**). When we stratified the performance across seven intrinsic functional networks[48], the sensory/motor regions demonstrated high accuracy with low diffusion times (mean correlation coefficients of visual/somatomotor = 0.775 / 0.886 at *t* = 1 and 0.881 / 0.931 at *t* = 10), whereas the transmodal regions of the frontoparietal and default mode networks showed low performance at low diffusion times but improved monotonically at higher diffusion times (frontoparietal/default mode = 0.721 / 0.672 at *t* = 1 and 0.845 / 0.837 at *t* = 10; **Fig. 1c**). The limbic region exhibited the lowest prediction performance compared with the other networks. These findings were consistent upon performing an analysis using the Desikan–Killiany-based sub-parcellation with 300 parcels [49,50] (**Supplement Fig. 1**).

**Fig. 1.**
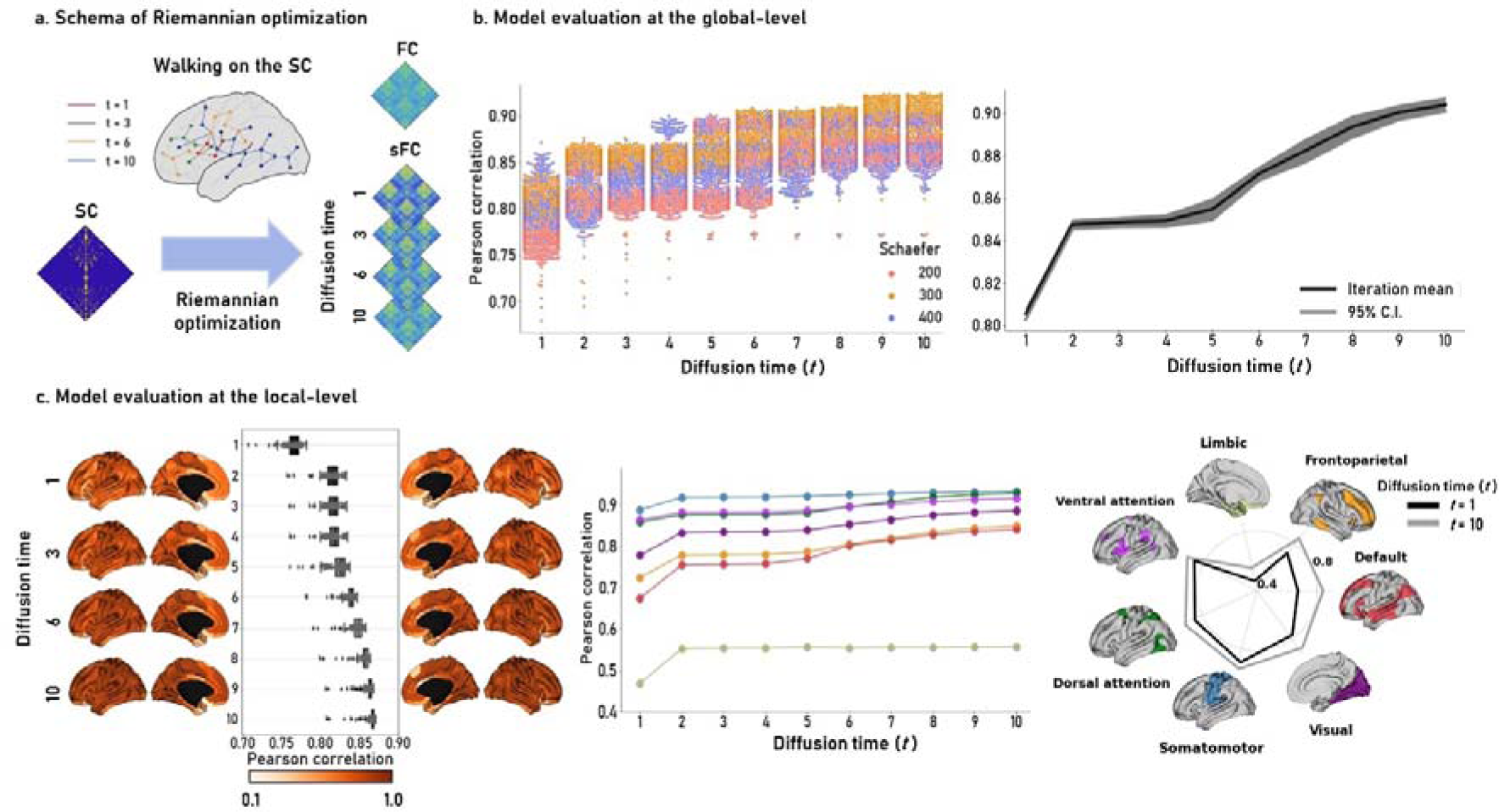
Structure-function coupling using Riemannian optimization. **(a)** Schema of the Riemannian optimization approach to simulate functional connectivity from the structural connectivity at different diffusion times (*t*). **(b)** We calculated the linear correlations between the actual and simulated functional connectivity across diffusion times for each individual (left). Each dot indicates each individual. We repeated the analysis 30 times with different training and test datasets for the Schaefer atlas with 300 parcels (right). The black line is the mean of 30 iterations, and the gray area indicates a 95% confidence interval (CI). **(c)** Regional prediction accuracy is shown on brain surfaces, and the box plots indicate the prediction accuracy of each brain region (left). The prediction performance was stratified according to the seven functional networks (middle and right). *Abbreviations:* SC, structural connectivity; FC, functional connectivity; sFC, simulated functional connectivity.

### Hierarchical organization of the functional gradients across diffusion times

To assess the cortical hierarchical organization of the functional connectome, we estimated the principal gradients from the simulated functional connectivity across the diffusion times (**Fig. 2a** and **Supplement Fig. 2**)[50,51]. The eigenvalue profile of the simulated gradients was highly similar to that estimated from the actual functional connectivity when the diffusion time was six (**Fig. 2b**). Specifically, the eigenvalue of the first simulated gradient was largely similar to the actual gradient (explained information = 46.0, 34.8, and 31.2% for simulated gradients at *t* = 1, 6, and 10; 34.8% for actual functional gradient). As the eigenvalues of the estimated gradients showed clear elbow between component number three and four, and the three gradients explained the information of the simulated functional connectivity with approximately 75, 71, and 68% at *t* = 1, 6, and 10, we thereafter considered three gradients. Similar to previous studies based on the HCP dataset[51], the first gradient differentiated sensory regions from transmodal regions, the second differentiated visual from somatomotor areas, and the third differentiated multiple demand networks from the rest of the brain. We summarized the distribution of the gradient values according to the seven functional networks across diffusion times. The SD of the distribution decreased at higher diffusion times, particularly in the frontoparietal and default mode networks (frontoparietal network:0.269, 0.216, and 0.209; default mode network:0.472, 0.410, and 0.356 at *t* = 1, 6, and 10; joy plot in **Fig. 2a**). To quantitatively evaluate the discrimination of gradient values between the sensory and transmodal regions, we calculated the distance of the distribution of gradient values between the visual/somatomotor and frontoparietal/default-mode networks. We confirmed that low-level sensory and higher-order transmodal regions were clearly differentiated at *t* = 6 (**Fig. 2c**). In the sensory areas, a noticeable change was observed between *t* = 1 and 2, and in the transmodal regions between *t* = 5 and 6. These findings indicated that sensory regions were well predicted with direct monosynaptic connections, whereas transmodal regions might be explained via indirect polysynaptic pathways. The changes in the second gradient across diffusion times showed large differences in the motor and visual areas, and the third gradient in the task-related areas (**Supplement Fig. 2**).

**Fig. 2.**
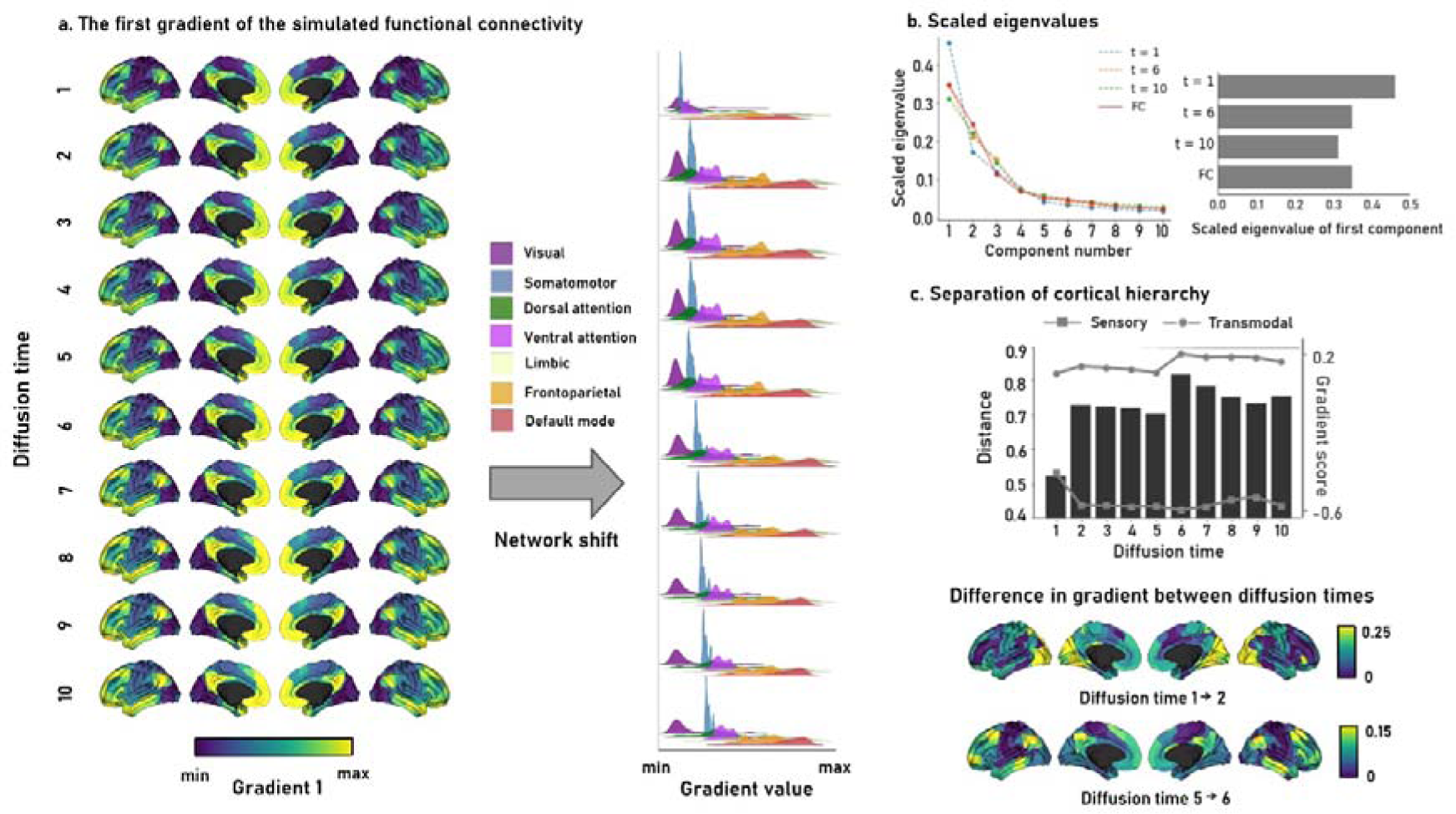
Differences in the cortical hierarchy of structure-function coupling across diffusion times. **(a)** The first gradient of the simulated functional connectivity is shown on brain surfaces across different diffusion times (left). The joy plots represent the distribution of the gradient values summarized based on the seven functional networks (right). **(b)** A scree plot describes information explained across the principal components of simulated and actual functional connectivity data. **(c)** We calculated the distance of the gradient value distribution between sensory and transmodal regions (top). The differences in gradient values that showed noticeable changes between diffusion times are shown on the brain surfaces (bottom).

### Associations with BMI

We associated the three simulated gradients with interindividual variations in BMI across diffusion times after controlling for age, sex, and head motion using the BrainStat toolbox (https://github.com/MICA-MNI/BrainStat)[52]. At diffusion times 5 and 6, we found significant associations in the frontoparietal and default mode regions, suggesting that variations in the BMI are related to the brain functions involved in higher-order cognitive control systems (**Fig. 3**). Indeed, the overall t-statistic values and number of significantly associated brain regions were particularly high at *t* = 6, which showed the clearest hierarchical differentiation.

**Fig. 3.**
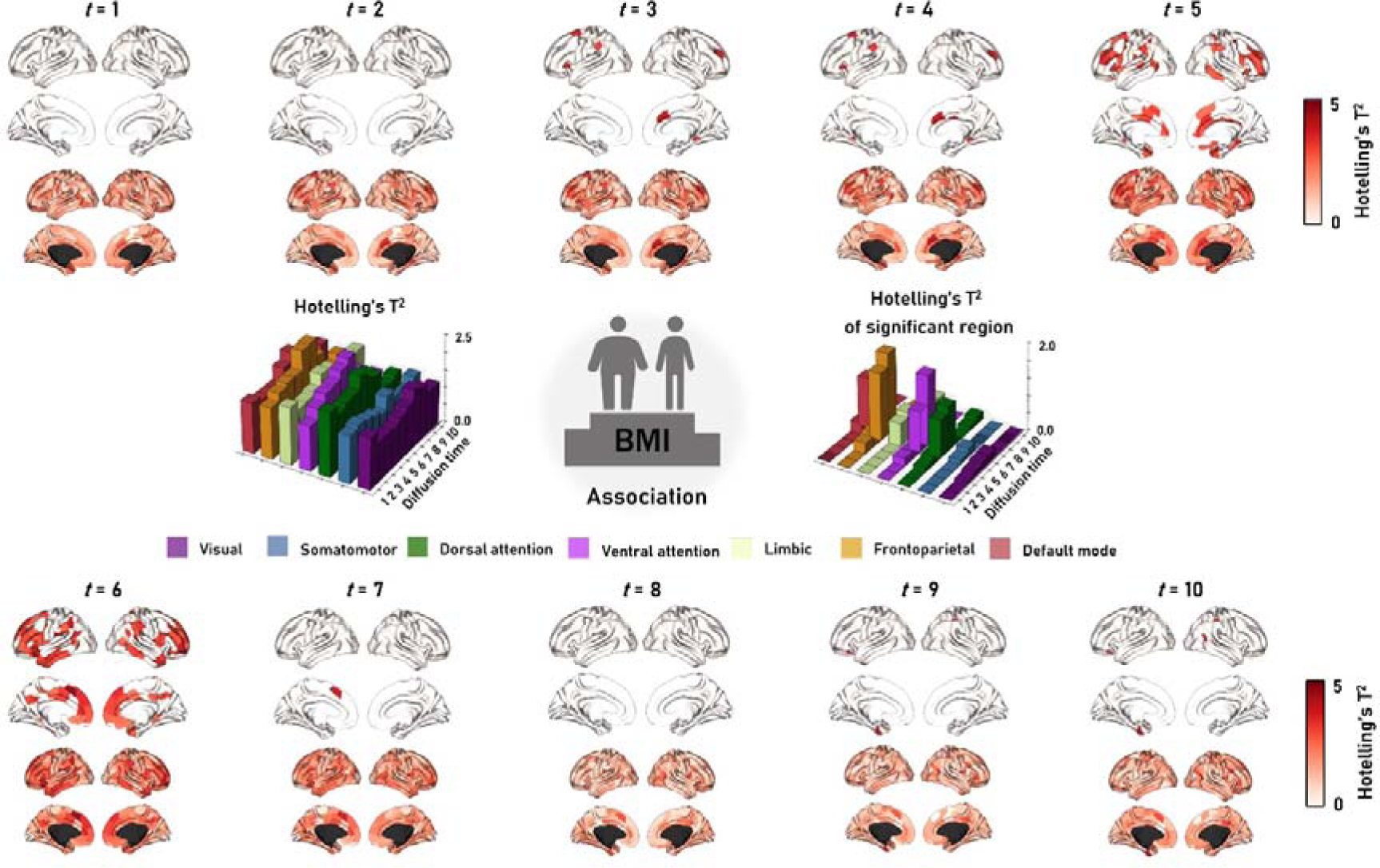
Associations between the simulated functional gradients and body mass index (BMI). Hotelling’s T^2^ statistics of the whole cortex and the regions that showed significant relations (FDR < 0.05) are shown on the brain surfaces across the diffusion times. The t-statistics are stratified according to seven functional networks and represented with bar plots. *Abbreviation:* FDR, false discovery rate.

### Cytoarchitectonic association analysis

To provide the biological underpinnings of BMI-related structure-function coupling, we estimated the microstructural gradient and moment features from BigBrain data, which are volumetrically reconstructed *post-mortem* data[53]. The strongest correlations were found at *t* = 6 (gradient: r = 0.393, p_spin-false discovery rate (FDR)_ < 0.001; mean: r = 0.316, p_spin-FDR_ = 0.029; SD: r = −0.410, p_spin-FDR_ < 0.001; skewness: r = 0.361, p_spin-FDR_ < 0.001; kurtosis: r = 0.362, p_spin-FDR_ < 0.001; **Fig. 4**). These findings suggest that alterations in structure-function coupling according to the BMI are associated with the brain microstructure, where the effects are dominant in transmodal areas with low laminar differentiation.

**Fig. 4.**
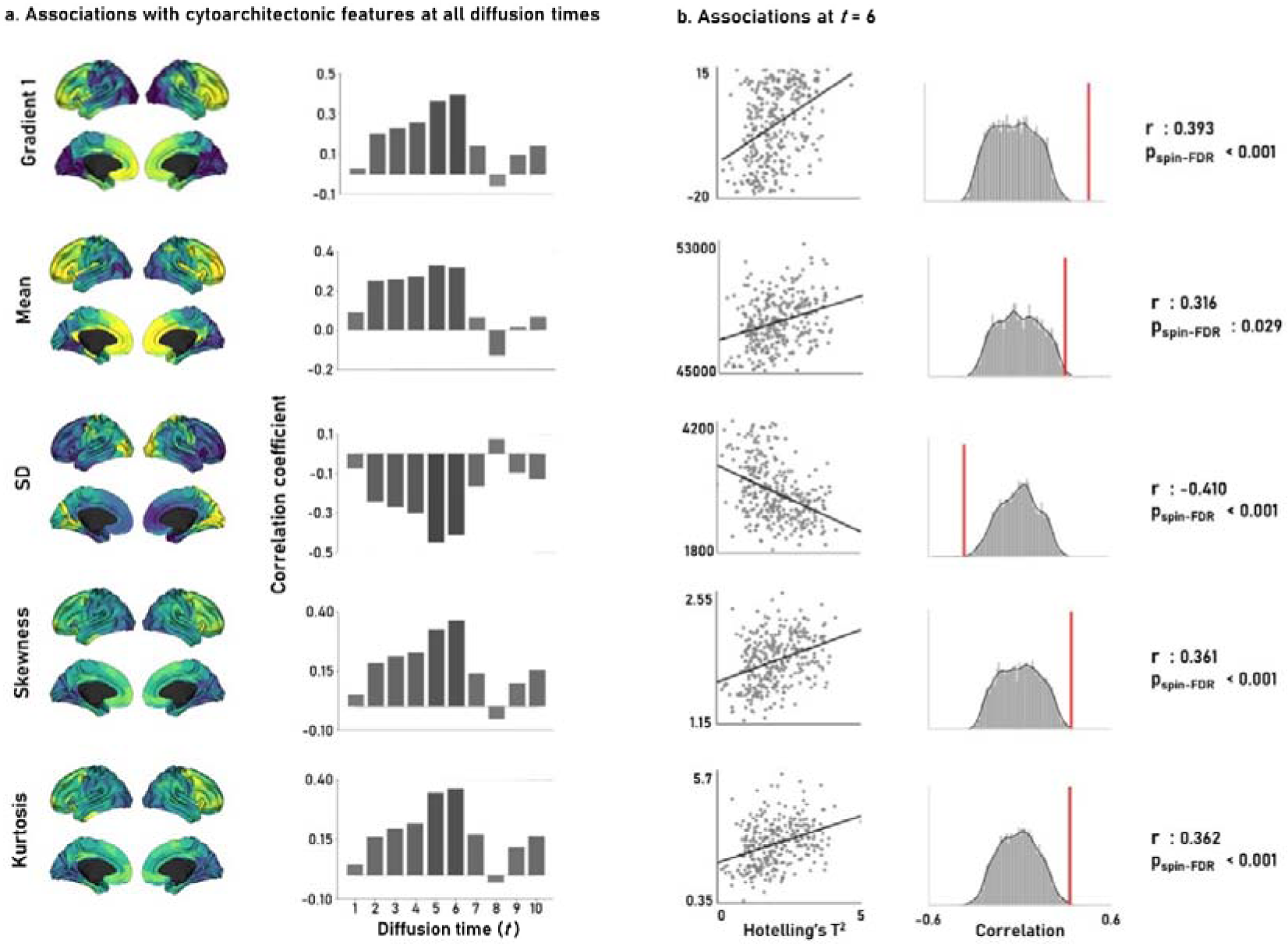
Cytoarchitectonic association analysis. **(a)** The distribution of cytoarchitectonic features (gradient, mean, SD, skewness, and kurtosis) are shown on the brain surfaces. The correlation coefficients between the cytoarchitectonic features and the structure-function coupling according to the BMI across diffusion times are shown with bar plots. **(b)** The correlation plots represent associations at diffusion time six. The distributions of correlation coefficients from 10,000 spin permutation tests are reported with histograms, and the actual correlation coefficients are represented with red lines. *Abbreviations:* SD, standard deviation; BMI, body mass index; FDR, false discovery rate.

### Transcriptomic association analysis

Additionally, we performed a transcriptomic association analysis with structure-function coupling according to the BMI to assess potential genetic relationships (**Fig. 5a**)[54,55]. Correlations with the gene expression data were high at diffusion times 5 and 6 (**Fig. 5b**). Cell type-specific expression analysis using the gene lists at *t* = 6 (**Supplementary Data**) showed enrichment of cells in the striatum, hypothalamus, and cortex (**Fig. 5c**)[56–60]. Consistent findings were obtained using the observed genes at diffusion time 5 (**Supplementary Fig. 3** and **Supplementary Data**).

**Fig. 5.**
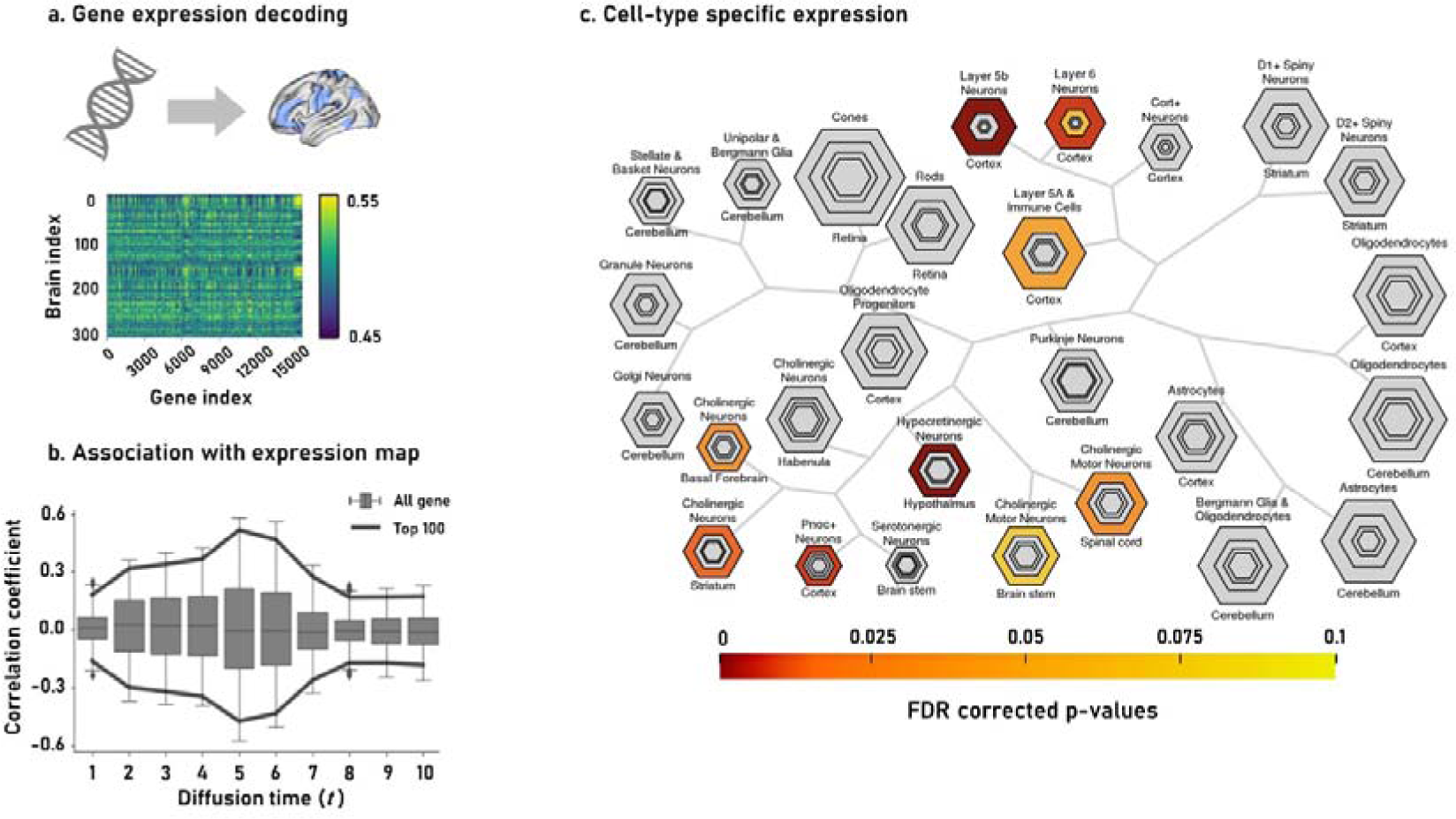
Transcriptomic association analysis. **(a)** The expression map of 15,677 genes in the brain regions. **(b)** The spatial correlations between the expression of each gene and the t-statistics of the BMI-related structure-function coupling are shown with box plots. Black lines represent the top 100 genes that showed negative and positive correlations. **(c)** Cell-type specific expression analysis identifies candidate cell populations. The hexagon size is scaled to the proportion of the gene lists, and varying stringencies for enrichment are represented by the size of the hexagons (specificity index threshold = 0.05, 0.01, 0.001, and 0.0001, respectively). Colors represent the FDR-corrected p-values. *Abbreviations:* BMI, body mass index; FDR, false discovery rate.

## DISCUSSION

Structure-function coupling is one of the key questions in neuroscience and may provide insights for understanding unseen neural mechanisms related to large-scale brain networks. In this study, we incorporated structural and functional information using a Riemannian optimization approach by varying the diffusion time parameters that reflect the implications of polysynaptic communication[38]. The simulation performance of the functional connectivity from the structural connectivity improved with increasing diffusion time, and the performance was remarkable in the transmodal regions, indicating that polysynaptic communication is required to simulate the higher-order functional dynamics of the transmodal systems. Moreover, we found that the hierarchical organization of the simulated functional gradients was most evident in the middle of the mono-and polysynaptic communications and that BMI was most significantly associated at these stages. BMI-related structure-function coupling was further associated with cytoarchitectonic and gene expression maps, suggesting the potential biological underpinnings of our findings.

Structure-function coupling has been actively studied in many previous studies, and higher-order transmodal regions require polysynaptic mechanisms, whereas low-level sensory systems can be modeled via relatively direct monosynaptic communication [27]–[34]. In this study, we adopted a Riemannian optimization approach to simulate the functional connectivity from structural information across different diffusion times to reflect polysynaptic communication[38]. This approach can perform the simulation at an individual participant level, enabling the investigation of the associations between structure and function coupling and behavioral or clinical traits at an individual level. We demonstrate the robustness of the model by changing the spatial granularities and parcellation schemes and applying a strict cross-validation framework. We consistently observed that the transmodal regions, including the frontoparietal and default-mode networks, were optimized when the diffusion time was high, indicating that these systems require multi-hop polysynaptic communication compared to low-level sensory systems. To systematically assess the topology of structurally governed functional networks, we estimated low-dimensional eigenvectors from the simulated functional connectivity. We found that the hierarchical organization differentiating the sensory and transmodal systems was notable in the middle of the diffusion times. The human brain has evolved to form a hierarchically organized cortical axis[61–64], and our findings may provide insights into random walk processes to describe the structurally governed functional dynamics along the cortex while forming cortical hierarchies.

Our study evaluated the associations between simulated functional gradients and inter-individual variations in BMI and highlighted significant effects on the frontoparietal and default mode networks at diffusion time, which revealed a notable cortical hierarchy. These findings suggest that variations in the BMI are associated with brain hierarchy. Indeed, previous studies complement our results in that individuals with high BMI show disruptions in the modular architecture and hierarchical organization of the brain, particularly in the transmodal regions[16,65]. Thus, variations in the BMI may be associated with synaptic communication along the cortical hierarchy, suggesting that obesity may reduce network communication efficiency among brain regions, resulting in delays in information transmission for the decision-making process. Furthermore, association analysis with cytoarchitectural measures revealed that BMI-related functional network alterations were associated with cortical layer-wise cell distribution. Our findings suggest that variations in the BMI may alter cell distribution within the cortical layers, which complement prior studies that demonstrated reduced neuronal density in the frontal and temporal regions in individuals with a high BMI[66].

Cell type-specific expression analysis provided further insights into BMI-related structure-function coupling. We observed associations among the striatum, hypothalamus, and cortical cells. The striatum plays an important role in reward processing[67], and the hypothalamus is involved in hormone and nutrient sensing, which maintains body weight, food intake, and energy expenditure[68]. Altered hypothalamic function is related to the dysfunction of the inhibitory circuit, inducing hedonic eating[69]. Indeed, previous studies have indicated that circuits involving the hypothalamus, striatum, and cortex are important for regulating metabolic homeostasis[70]. Although direct relationships between gene expression and macroscale structure-function coupling should be investigated in more depth, our cell type-specific expression analysis demonstrated the potential for constructing a consolidated framework linking BMI-related macroscale brain network reconfiguration and microscale perspectives.

In summary, we observed strong associations between BMI and structure-function coupling when the cortical hierarchy was clearly differentiated, and these associations were further related to brain microstructure and gene expression. Our findings provide insights into the polysynaptic communication mechanisms of BMI-related structure-function coupling in large-scale brain networks.

## METHODS

### Participants

We obtained T1- and T2-weighted structural, diffusion, and resting-state functional MRI data from the S1200 release of the Human Connectome Project (HCP) database (http://www.humanconnectome.org/)[47]. Among the 1,206 participants, we excluded those who were genetically related (i.e., twins), and had a family history of mental illness, history of drug ingestion, and incomplete multimodal imaging data. Finally, 290 participants (mean ± SD age = 28.3 ± 3.9 years; 51.3 % female) were included in our analysis. The mean and SD of BMI were 26.06 ± 4.85 kg/m^2^, and the rangeCwas between 17.01 and 42.91 kg/m^2^.

### MRI acquisition

MRI images were obtained using a Siemens Skyra 3T scanner at the University of Washington. The T1-weighted images were acquired using a magnetization-prepared rapid gradient echo (MPRAGE) sequence (repetition time [TR]C=C2400Cms; echo time [TE]C=C2.14Cms; field of view [FOV]C=C224C×C224Cmm^2^; voxel sizeC=C0.7Cmm^2^; number of slicesC=C256), and the T2-weighted MRI data were scanned using a T2-SPACE sequence, which had the same parameters as the T1-weighted data but different TR (3,200 ms) and TE (565 ms). Diffusion MRI data were obtained using a spin-echo echo-planar imaging (SE-EPI) sequence (TR = 8700 ms, TE = 90 ms, flip angle = 90°, voxel size = 2 mm^3^, 70 slices, FOV = 192 × 192 mm^2^, matrix size = 96 × 96 × 70, b-value = 1000 s/mm^2^, 63 diffusion directions, six b0 images). Resting-state functional MRI data were acquired using a gradient-echo EPI sequence (TRC=C720Cms, TEC=C33.1Cms; FOVC=C208C×C180Cmm^2^, voxel sizeC=C2Cmm^3^, number of slicesC=C72, and number of volumesC=C1,200). During the functional MRI scan, participants were instructed to keep their eyes open while looking at a fixed cross. Functional MRI consisted of two sessions, each containing data in a phase-encoded direction from left-to-right and right-to-left, providing up to four time series per participant.

### MRI data preprocessing

The HCP database provides minimally preprocessed data using FSL, FreeSurfer, and Workbench[71–73]. Briefly, the T1- and T2-weighted images were corrected for gradient nonlinearity and b0 distortions, and co-registered using a rigid-body transformation. Bias field correction was performed based on the inverse intensities from T1- and T2-weighting. The processed data were nonlinearly registered onto the Montreal Neurological Institute (MNI152) standard space, and the pial and white surfaces were generated by following the boundaries between the different tissues[74–76]. The mid-thickness surface was generated by averaging the pial and white surfaces and was used to create an inflated surface. The spherical surface was registered to the Conte69 template using MSMAll, and 164k vertices were downsampled to a 32k vertex mesh[77,78]. Diffusion MRI data were corrected for susceptibility distortions, head motion, and eddy currents [44]. Resting-state functional MRI data were corrected for EPI distortion and head motion. Data were registered to the T1-weighted image and subsequently to the MNI152 space, and magnetic field bias correction, skull removal, and intensity normalization were performed. Noise components resulting from head movement, white matter, heartbeat, and arterial- and aortic-related contributions were removed using the FMRIB’s ICA-based X-noiseifier (ICA-FIX)[79]. Using a cortical ribbon-constrained volume-to-surface mapping algorithm, the time-series data were mapped to a standard gray ordinate space.

### Structural and functional connectivity construction

Structural connectomes were generated from preprocessed diffusion MRI data using MRtrix3[80]. Anatomically constrained tractography was performed using T1-weighted image-driven tissue types, including the cortical and subcortical gray matter, white matter, and cerebrospinal fluid[81]. After co-registering the T1-weighted and diffusion MRI data using boundary-based registration, we applied a transformation to the tissue types to align them in the native diffusion MRI space. We estimated multishell and multitissue response functions and performed constrained spherical deconvolution and intensity normalization[82]. Tractograms were generated by seeding from all white matter voxels using a probabilistic approach [83] with 40 million streamlines, a maximum tract length of 250, and a fractional anisotropy cutoff of 0.06. We subsequently applied spherical deconvolution informed filtering of tractograms (SIFT2) to reconstruct the streamlines weighted by cross-section multipliers[84]. Finally, the structural connectivity matrix was constructed using the Yeo seven-network-based Schaefer atlas with 200, 300, and 400 parcels, [48,85] as well as the Desikan–Killiany-based sub-parcellation with 300 parcels [49,50]and log-transformed. Functional connectomes were constructed by calculating linear correlations of functional time series between two different regions defined using the Schaefer atlas with 200, 300, and 400 parcels [48,85] and Desikan–Killiany-based sub-parcellation with 300 parcels[49,50]. The correlation coefficients were Fisher’s r-to-z-transformed to render the data more normally distributed[86].

### Riemannian optimization for structure-function coupling

To assess structure-function coupling, we adopted the Riemannian optimization approach[38]. Briefly, it generates low-dimensional eigenvectors (i.e., diffusion maps) from the structural connectivity matrix by applying a nonlinear dimensionality reduction technique (i.e., diffusion map embedding)[87]. The diffusion maps are controlled by the diffusion time parameter, *t*, where a higher *t* embeds the data more closely in the low-dimensional manifold space. By varying the diffusion time parameter between *t* = 1 and 10, we applied a kernel fusion approach to simulate functional connectivity so that the functional connectivity and diffusion maps had minimum differences. Specifically, the algorithm determines a transformation matrix that rotates the diffusion maps to reconstruct the functional connectivity. Thus, the rotation matrix may indicate optimal paths for propagating information between different brain regions. We performed a prediction analysis with fivefold cross-validation and repeated the analysis 30 times with different training and test datasets to mitigate potential subject selection bias. Model performance was evaluated by calculating the linear correlation between the elements of the actual and simulated functional connectivity matrices. At the regional level, we calculated the linear correlations between actual and simulated functional connectivity for each brain region. We stratified the correlation coefficients according to seven intrinsic functional communities [48] to evaluate the results in terms of functionally differentiated networks.

### Simulated functional gradients across diffusion times

We estimated low-dimensional representations of functional connectivity (i.e., gradients) to assess differences in the cortical hierarchy across different diffusion times. Using the BrainSpace toolbox (https://github.com/MICA-MNI/BrainSpace)[50], we applied diffusion map embedding [87] to the affinity matrix, which was constructed by applying a normalized angle kernel to the group-averaged connectivity matrix. Individual gradients were then estimated and aligned to the group template gradient using Procrustes analysis[88]. After normalizing the gradient values between −1 and 1 using a min–max scaling at each diffusion time, we evaluated the shifts in the distribution of the values of the first functional gradient for each functional network[48]. Additionally, we calculated the distance of the gradient value distributions between the sensory (visual and somatomotor) and transmodal networks (frontoparietal and default modes) at each diffusion time point to assess the hierarchical differentiation of the brain.

### Associations between structure-function coupling and BMI

We used a standard general linear model to assess the associations between interindividual variations in the BMI and structure-function coupling at different diffusion times using the BrainStat toolbox (https://github.com/MICA-MNI/BrainStat)[52]. After controlling for age, sex, and head movement, defined using framewise displacement, we estimated the associations between the BMI and the three simulated functional gradients. The inference was based on Hotelling’s t-square statistics, and multiple comparisons across brain regions were corrected using FDR[89].

### Cytoarchitectonic and transcriptomic association analyses

To elucidate the biological underpinnings of the association between BMI and structure-function coupling measures, we performed cytoarchitectonic and transcriptomic association analyses. Cytoarchitectural features were calculated from the BigBrain dataset, an ultra-high-resolution, three-dimensional volumetric reconstruction of a *post-mortem* Merker-stained and sliced human brain from a 65-year-old male (https://bigbrain.loris.ca/main.php)[53]. Data were mapped onto a Schaefer atlas with 300 parcels[85]. We first generated 14 equivolumetric cortical surfaces within the cortex (https://github.com/caseypaquola/BigBrainWarp) and sampled intensity values along these surfaces. We then constructed a microstructural profile covariance matrix based on the linear correlations of the cortical depth-dependent intensity profiles between different brain regions, controlling for the average whole-cortex intensity profile[90]. The matrix was thresholded at zero and log-transformed, and a microstructural gradient was generated using BrainSpace (https://github.com/MICA-MNI/BrainSpace)[50]. An affinity matrix was constructed using a normalized angle kernel with the top 10% entries for each parcel and we applied diffusion map embedding to generate a microstructural gradient [87]. Next, we calculated four moment features (mean, SD, skewness, and kurtosis) from the microstructural profile of each brain region. Mean and SD represent the overall intensity distribution across the cortical layers, skewness represents the shift in intensity values towards the supragranular layer (positive skewness) or flat distribution (negative skewness), and kurtosis indicates whether the tail of the intensity distribution contains extreme intensity values[41]. We associated the t-statistics of BMI and structure-function coupling relationships with the cytoarchitectonic features by calculating linear correlations with 1,000 spin permutation tests to account for spatial autocorrelations[91]. Multiple comparisons across features were corrected using FDR[89].

Transcriptomic association analysis was performed using the Abagen toolbox (https://abagen.readthedocs.io/en/stable/)[54,55]. The toolbox processed a dataset containing microarray expression data collected from six *post-mortem* human brains obtained from the Allen Human Brain Atlas[92]. We mapped the expression data onto parcels, and the missing data were interpolated by assigning the expression of the nearest tissue sample. As four of the six donors provided expression maps only in the left hemisphere, the expression data were mirrored from the left to right hemispheres. We used the differential stability method to select probes with multiple probe indexed expression. It computes the Spearman correlation of the expression data for every pair of donors, and the probe with the highest correlation is retained. The expression data were normalized using a scaled robust sigmoid function to mitigate potential differences in the expression values among donors. Finally, a gene expression map was constructed by averaging the expression data from six donors. We subsequently calculated the linear correlations between each gene expression data point and the t-statistics of BMI and structure-function coupling according to the diffusion times. In addition, we conducted cell-type-specific expression analysis (http://genetics.wustl.edu/jdlab/csea-tool-2/) using the genes that demonstrated an absolute correlation coefficient > 0.45 and passed for FDR < 0.05[93].

## DATA AVAILABILITY

Imaging and phenotypic data were provided in part by the Human Connectome Project (HCP; http://www.humanconnectome.org/).

## CODE AVAILABILITY

The codes for the Riemannian optimization are available at https://github.com/MICA-MNI/micaopen/tree/master/sf_prediction, codes for gradient generation at https://github.com/MICA-MNI/BrainSpace, and codes for statistical analyses are available at https://github.com/MICA-MNI/BrainStat.

## Supplementary information

**Supplementary Fig. 1.**
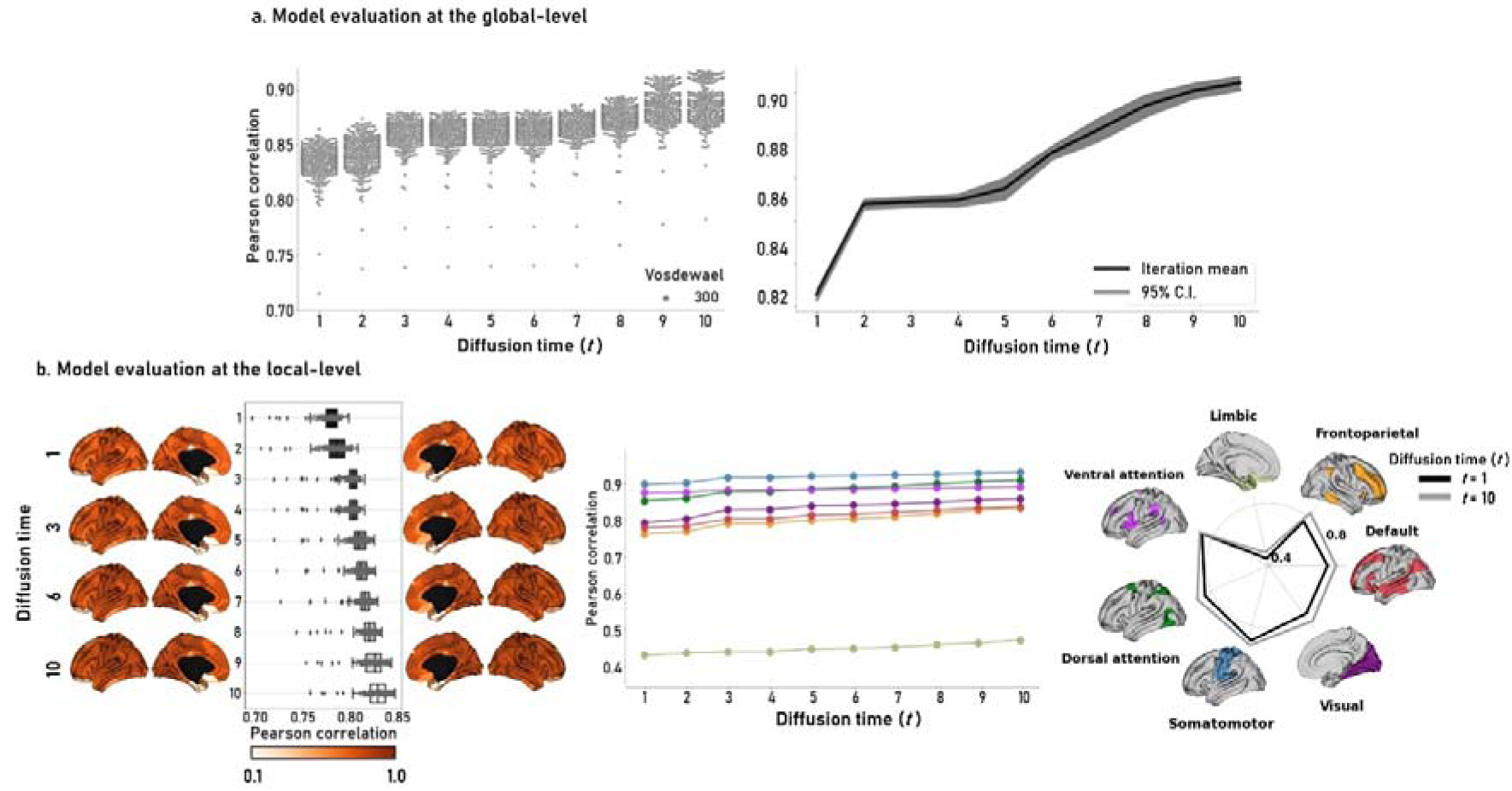
Structure-function coupling using Riemannian optimization based on the Desikan–Killiany-based sub-parcellation with 300 parcels. **(a)** We calculated linear correlations between the actual and simulated functional connectivity across diffusion times for each individual (left). Each dot indicates each individual. We repeated the analysis 30 times with different training and test datasets for the Schaefer atlas with 300 parcels (right). The black line is the mean of 30 iterations, and the gray area indicates a 95% confidence interval (CI). **(b)** Regional prediction accuracy is demonstrated on the brain surfaces, and the box plots indicate the prediction accuracy of each brain region (left). The prediction performance was stratified according to the seven functional networks (middle and right).

**Supplementary Fig. 2.**
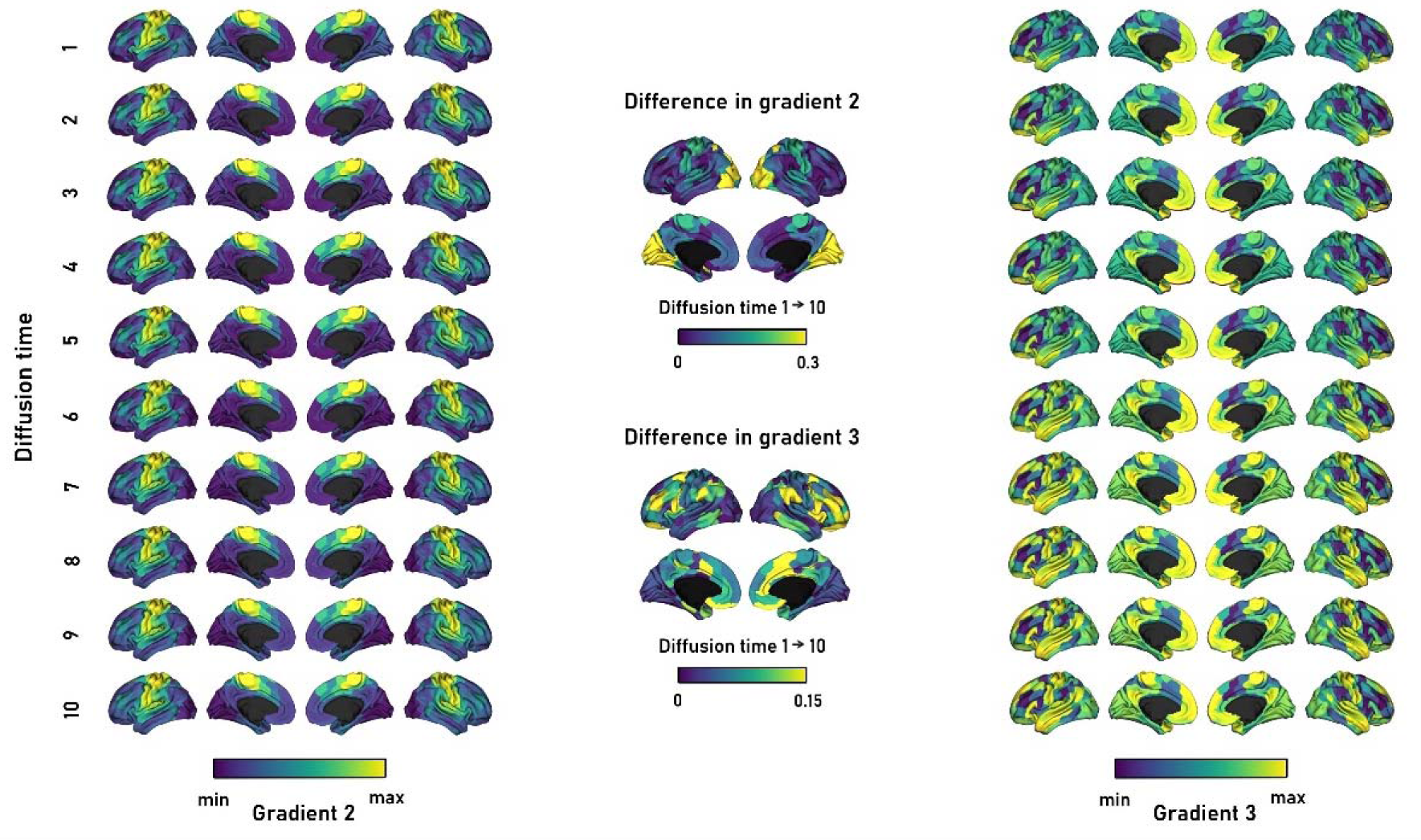
Hierarchical organization of the second and third simulated functional gradients across diffusion times. The second and third gradients of simulated functional connectivity are shown (left and right). The differences in the gradient values between *t* = 1 and 10 are shown on the brain surfaces (middle).

**Supplementary Fig. 3.**
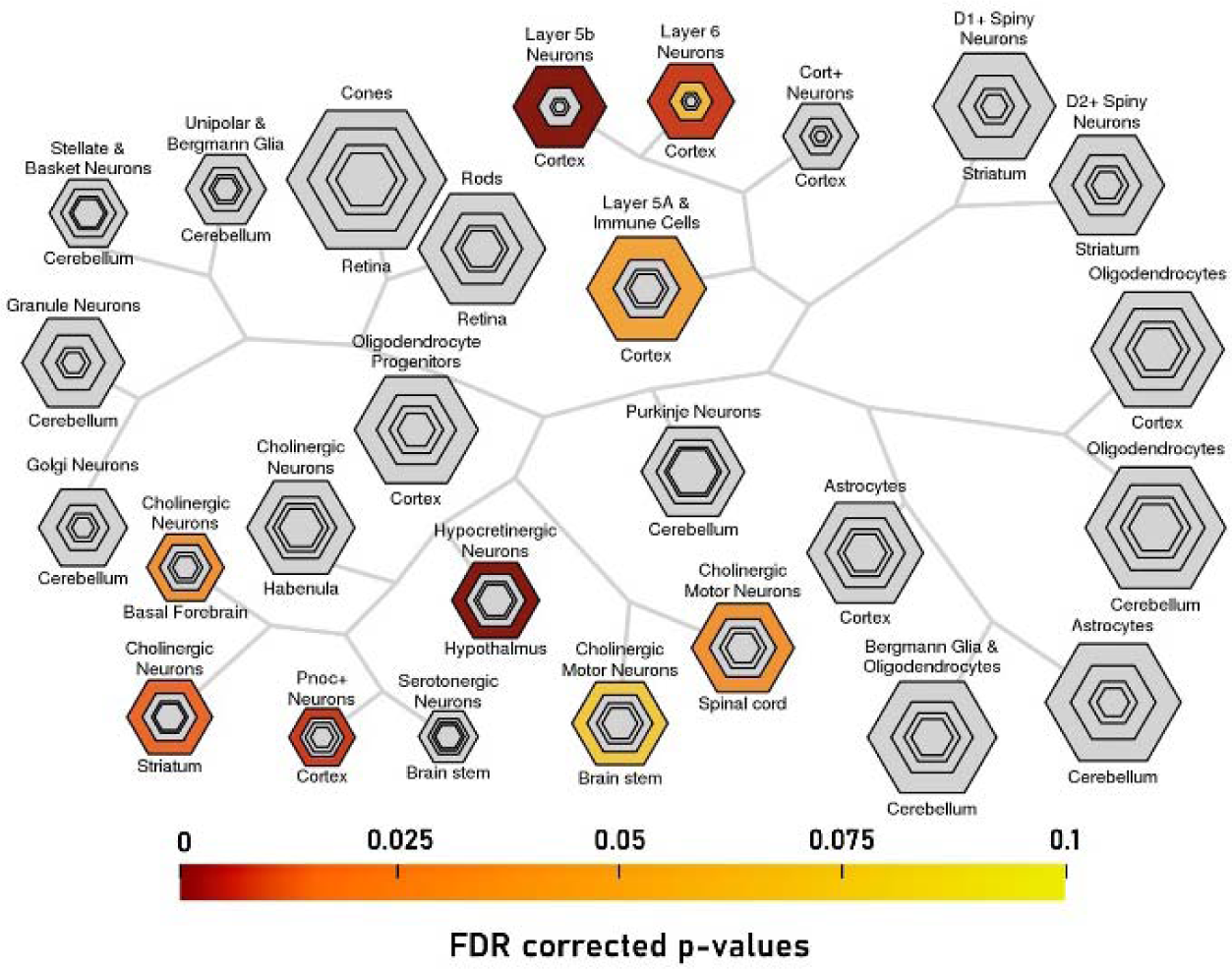
Transcriptomic association analysis. The correlation coefficient between the t-statistic map at diffusion time 5 and the gene expression was calculated, and we performed cell-type specific expression analysis.

## Supplementary Data

**A total of 201 genes that were associated with BMI-related structure-function coupling at diffusion time 5**

[′ACAN′, ′ACOX3′, ′ACSL6′, ′ADAMTS3′, ′ADM′, ′ANKRD55′, ′AP3S1′, ′ARHGAP28′, ′ARHGAP33′, ′ARHGDIG′, ′ASS1′, ′ATP2B3′, ′ATP9A′, ′B3GAT1′, ′B4GALT2′, ′BAIAP3′, ′BATF3′, ′BCRP2′, ′C11orf97′, ′C17orf67′, ′C1S′, ′C2CD4C′, ′C2orf80′, ′C4orf33′, ′C8orf46′, ′CA10′, ′CCDC3′, ′CCDC58′, ′CCNYL1′, ′CDH10′, ′CDH7′, ′CDH8′, ′CDKN2D′, ′CEP170B′, ′CHCHD6′, ′CLGN′, ′CNIH2′, ′CRIP2′, ′CTC1′, ′CTXN1′, ′CUX1′, ′CXorf57′, ′CYP46A1′, ′DCAF11′, ′DDA1′, ′DDN′, ′DDRGK1′, ′DOK6′, ′DPF1′, ′DPP10-AS1′, ′DPYSL3′, ′DUSP3′, ′DYRK2′, ′DZIP3′, ′E2F3′, ′EEPD1′, ′EFCAB1′, ′EFHC2′, ′EFNB2′, ′EFNB3′, ′ENOX1′, ′EPHA10′, ′EPOP′, ′ERFE′, ′EYA4′, ′F12′, ′FAM102B′, ′FAM110C′, ′FASTKD1′, ′FIGNL2′, ′FREM3′, ′FSTL1′, ′FXYD6′, ′FZR1′, ′GABRB1′, ′GAS2′, ′GGN′, ′GLRA3′, ′GMFB′, ′GSG1′, ′GSS′, ′GSTM3′, ′GULP1′, ′HDAC9′, ′HDC′, ′HES4′, ′HRH1′, ′HS3ST1′, ′HSD11B1L′, ′HTR7P1′, ′IFT22′, ′IP6K2′, ′ITGA11′, ′JPH3′, ′KCNA5′, ′KCNC4′, ′KCNG3′, ′KCNMB4′, ′KCTD15′, ′KIF21B′, ′KIRREL2′, ′L3MBTL4′, ′LAG3′, ′LCA5′, ′LINC00484′, ′LINC01102′, ′LINC01137′, ′LINC01197′, ′LINC02217′, ′LOC100288911′, ′LOC100294145′, ′LOC440934′, ′LOC728392′, ′LRRC2′, ′LRRC49′, ′LY6H′, ′LYPD8′, ′MBOAT7′, ′MC1R′, ′MEA1′, ′MOB1B′, ′MORN4′, ′MRAS′, ′MSANTD1′, ′MTA3′, ′MUM1L1′, ′NAALAD2′, ′NANOS1′, ′NEB′, ′NEUROD6′, ′NEXN′, ′NLE1′, ′NOL4′, ′NOV′, ′NT5DC2′, ′NT5DC3′, ′NUDT14′, ′OPRM1′, ′PCBD1′, ′PCNT′, ′PECR′, ′PI4K2A′, ′PLCD4′, ′PLPPR3′, ′PLXNA1′, ′PNCK′, ′PPP4R4′, ′PRKCD′, ′PRR16′, ′PTPRF′, ′PUSL1′, ′PYGL′, ′QRFPR′, ′RAB36′, ′RASAL1′, ′RASSF4′, ′RBP4′, ′RGS4′, ′RNF150′, ′RRAS2′, ′RSPO2′, ′SCARA5′, ′SCN3B′, ′SH2D5′, ′SHISA9′, ′SHISAL1′, ′SIDT1′, ′SLC25A23′, ′SLC26A4-AS1′, ′SLC46A3′, ′SMIM10L2A′, ′SNHG8′, ′SNX7′, ′SPRN′, ′ST3GAL6′, ′ST6GALNAC5′, ′STX1A′, ′SUSD1′, ′SVOP′, ′SYNE4′, ′TBC1D24′, ′TCEA3′, ′TFPT′, ′TM2D3′, ′TMEM108′, ′TMEM130′, ′TMEM145′, ′TMEM150C′, ′TOM1L1′, ′TRPC3′, ′TSPAN33′, ′TTPAL′, ′TXN′, ′VAV3′, ′VIT′, ′WDR31′, ′WDR86′, ′WNT10B′, ′WTIP′, ′XYLT1′, ′ZNF350′]

**A total of 230 genes that associated with BMI-related structure-function coupling at diffusion time 6**

[′ACOX3′, ′ACTC1′, ′ADAMTS3′, ′ADCY2′, ′ADCY7′, ′ADTRP′, ′AGPAT2′, ′AMIGO2′, ′ANKRD55′, ′ARHGAP28′, ′ARHGDIG′, ′ARL5A′, ′ARPP19′, ′ASS1′, ′ATP9A′, ′B3GAT1′, ′BAIAP2L2′, ′BAIAP3′, ′BATF3′, ′BTN2A2′, ′C17orf67′, ′C1S′, ′C2CD4C′, ′C4orf33′, ′C8orf46′, ′CA10′, ′CACNG3′, ′CAMK1G′, ′CCDC136′, ′CCDC3′, ′CCDC58′, ′CCDC68′, ′CCK′, ′CCKBR′, ′CCNYL1′, ′CD83′, ′CDH10′, ′CDH8′, ′CDKN2D′, ′CEP170B′, ′CHCHD3′, ′CHCHD6′, ′CHRDL1′, ′CHRM2′, ′CIB1′, ′CLGN′, ′CLIP4′, ′CMPK1′, ′CNTLN′, ′CRIP2′, ′CRYM′, ′CTC1′, ′CTXN1′, ′CUX1′, ′CYB561′, ′CYP46A1′, ′DCAF11′, ′DCP2′, ′DDRGK1′, ′DGKB′, ′DLEU7′, ′DLK2′, ′DOC2A′, ′DOK6′, ′DPF1′, ′DPP10-AS1′, ′DUSP3′, ′DYRK2′, ′DZIP3′, ′E2F3′, ′EEPD1′, ′EFCAB1′, ′EFNB2′, ′ENOX1′, ′EPHA10′, ′EPOP′, ′FADS3′, ′FAM102B′, ′FAM110C′, ′FAM13B′, ′FASTKD1′, ′FKBP5′, ′FREM3′, ′FRMD4A′, ′FSTL1′, ′FXYD6′, ′GAP43′, ′GAS2′, ′GAS6′, ′GGN′, ′GIT2′, ′GLRA3′, ′GMFB′, ′GRASP′, ′GSG1′, ′GSS′, ′GUCA2B′, ′GULP1′, ′HDAC9′, ′HDC′, ′HERC3′, ′HRH1′, ′HS3ST1′, ′HSD11B1L′, ′HSPA4L′, ′HTR7P1′, ′IP6K2′, ′JMJD1C-AS1′, ′KCNA5′, ′KCNC4′, ′KCNMB4′, ′KIF21B′, ′KIRREL2′, ′KLF6′, ′KREMEN1′, ′L3MBTL4′, ′LAMB1′, ′LCA5′, ′LINC-PINT′, ′LINC00484′, ′LINC02217′, ′LOC440934′, ′LRRC2′, ′LRRC49′, ′LRRC73′, ′LY6H′, ′LY86-AS1′, ′LYPD8′, ′MAPK11′, ′MC1R′, ′MCUB′, ′MEA1′, ′MEIS3′, ′MFSD9′, ′MKL2′, ′MOB1B′, ′MORN4′, ′MRAS′, ′MSANTD1′, ′MTA3′, ′MTCH1′, ′MUM1L1′, ′MYDGF′, ′NFKBIE′, ′NIPA1′, ′NOL4′, ′NOV′, ′NT5DC2′, ′NT5DC3′, ′NUDT14′, ′NUDT4′, ′OLFM3′, ′OPRM1′, ′PCBD1′, ′PCTP′, ′PDZD8′, ′PECR′, ′PI4K2A′, ′PIK3CD′, ′PLCD4′, ′PLXNA1′, ′PPEF1′, ′PRKCD′, ′PRNP′, ′PTPRF′, ′QRFPR′, ′RASAL1′, ′RASL11B′, ′RASSF4′, ′RBP4′, ′RETREG1′, ′RGS4′, ′RRM2B′, ′RSPO2′, ′RTL8C′, ′RTP1′, ′RUNDC3A′, ′SCARA5′, ′SCLT1′, ′SECISBP2′, ′SFTPD′, ′SH2D5′, ′SHISAL1′, ′SLC26A4-AS1′, ′SLC35A2′, ′SLC46A3′, ′SLC4A9′, ′SMIM10L2A′, ′SMIM10L2B′, ′SPICE1′, ′SPINT2′, ′SPON2′, ′SPPL3′, ′SPRN′, ′SRFBP1′, ′ST3GAL6′, ′ST6GALNAC5′, ′STMN1′, ′STX12′, ′STX1A′, ′SVOP′, ′SYNE4′, ′TBC1D24′, ′TIAM1′, ′TM2D3′, ′TMEM108′, ′TMEM130′, ′TMEM150C′, ′TOM1L1′, ′TPD52′, ′TPST1′, ′TRIM27′, ′TSPAN33′, ′TTC21B′, ′TTPAL′, ′TUBB2A′, ′TUBB6′, ′UBE2G1′, ′UBE2Z′, ′VAV3′, ′VIT′, ′VSTM2L′, ′WDR31′, ′WDR86′, ′WNT10B′, ′WTIP′, ′XYLT1′, ′YTHDF2′, ′YWHAH′, ′ZBTB44′]

## REFERENCES

1. Ng M, Fleming T, Robinson M, Thomson B, Graetz N, Margono C, et al. Global, regional, and national prevalence of overweight and obesity in children and adults during 1980-2013: A systematic analysis for the Global Burden of Disease Study 2013. The Lancet. 2014;384: 766–781. doi:10.1016/S0140-6736(14)60460-8

2. Malik VS, Willett WC, Hu FB. Global obesity: trends, risk factors and policy implications. Nat Rev Endocrinol. 2013;9: 13–27. doi:10.1038/nrendo.2012.199

3. Gosmanov AR, Smiley D, Robalino G, Siqueira JM, Peng L, Kitabchi AE, et al. Effects of intravenous glucose load on insulin secretion in patients with ketosis-prone diabetes during near-normoglycemia remission. Diabetes Care. 2010;33: 854–860. doi:10.2337/dc09-1687

4. Studies Collaboration P. Body-mass index and cause-specifi c mortality in 900 000 adults: collaborative analyses of 57 prospective studies Prospective Studies Collaboration*. The Lancet. 373: 1083–1096. doi:10.1016/S0140

5. Vgontzas AN, Bixler EO, Chrousos GP. Obesity-related sleepiness and fatigue: The role of the stress system and cytokines. Ann N Y Acad Sci. 2006;1083: 329–344. doi:10.1196/ANNALS.1367.023

6. Nedunchezhiyan U, Varughese I, Sun ARJ, Wu X, Crawford R, Prasadam I. Obesity, Inflammation, and Immune System in Osteoarthritis. Front Immunol. 2022;13. doi:10.3389/FIMMU.2022.907750/FULL

7. Smith E, Hay P, Campbell L, Trollor JN. A review of the association between obesity and cognitive function across the lifespan: implications for novel approaches to prevention and treatment. Obesity Reviews. 2011;12: 740–755. doi:https://doi.org/10.1111/j.1467-789X.2011.00920.x

8. Charidimou A, Gang Q, Werring DJ. Sporadic cerebral amyloid angiopathy revisited: recent insights into pathophysiology and clinical spectrum. J Neurol Neurosurg Psychiatry. 2012;83: 124–137. doi:10.1136/JNNP-2011-301308

9. Gauthier MS, Ruderman NB. Adipose tissue inflammation and insulin resistance: all obese humans are not created equal. Biochem J. 2010;430. doi:10.1042/BJ20101062

10. Mariman ECM, Wang P. Adipocyte extracellular matrix composition, dynamics and role in obesity. Cell Mol Life Sci. 2010;67: 1277–1292. doi:10.1007/S00018-010-0263-4

11. Xu J, Li Y, Lin H, Sinha R, Potenza MN. Body mass index correlates negatively with white matter integrity in the fornix and corpus callosum: A diffusion tensor imaging study. Hum Brain Mapp. 2013;34: 1044–1052. doi:10.1002/hbm.21491

12. Carbine KA, Duraccio KM, Hedges-Muncy A, Barnett KA, Kirwan CB, Jensen CD. White matter integrity disparities between normal-weight and overweight/obese adolescents: an automated fiber quantification tractography study. Brain Imaging Behav. 2020;14: 308–319. doi:10.1007/s11682-019-00036-4

13. Taki Y, Kinomura S, Sato K, Inoue K, Goto R, Okada K, et al. Relationship Between Body Mass Index and Gray Matter Volume in 1,428 Healthy Individuals. Obesity. 2008;16: 119–124. doi:10.1038/OBY.2007.4

14. Medic N, Ziauddeen H, Ersche KD, Farooqi IS, Bullmore ET, Nathan PJ, et al. Increased body mass index is associated with specific regional alterations in brain structure. Int J Obes. 2016;40: 1177–1182. doi:10.1038/ijo.2016.42

15. Medic N, Kochunov P, Ziauddeen H, Ersche KD, Nathan PJ, Ronan L, et al. BMI-related cortical morphometry changes are associated with altered white matter structure. Int J Obes. 2019;43: 523–532. doi:10.1038/s41366-018-0269-9

16. Park B yong, Park H, Morys F, Kim M, Byeon K, Lee H, et al. Inter-individual body mass variations relate to fractionated functional brain hierarchies. Communications Biology 2021 4:1. 2021;4: 1–12. doi:10.1038/s42003-021-02268-x

17. Kullmann S, Heni M, Veit R, Ketterer C, Schick F, Häring HU, et al. The obese brain: Association of body mass index and insulin sensitivity with resting state network functional connectivity. Hum Brain Mapp. 2012;33: 1052. doi:10.1002/HBM.21268

18. Spiegelman BM, Flier JS. Obesity and the Regulation of Energy Balance. Cell. 2001;104: 531–543. doi:10.1016/S0092-8674(01)00240-9

19. Rubinov M, Sporns O. Complex network measures of brain connectivity: Uses and interpretations. Neuroimage. 2010;52: 1059–1069. doi:10.1016/J.NEUROIMAGE.2009.10.003

20. Bullmore E, Sporns O. Complex brain networks: graph theoretical analysis of structural and functional systems. Nature Reviews Neuroscience 2009 10:3. 2009;10: 186–198. doi:10.1038/nrn2575

21. Baum GL, Cui Z, Roalf DR, Ciric R, Betzel RF, Larsen B, et al. Development of structure–function coupling in human brain networks during youth. Proc Natl Acad Sci U S A. 2020;117: 771–778. doi:10.1073/PNAS.1912034117/SUPPL_FILE/PNAS.1912034117.SAPP.PDF

22. Hermundstad AM, Bassett DS, Brown KS, Aminoff EM, Clewett D, Freeman S, et al. Structural foundations of resting-state and task-based functional connectivity in the human brain. Proc Natl Acad Sci U S A. 2013;110: 6169–6174. doi:10.1073/PNAS.1219562110/SUPPL_FILE/SAPP.PDF

23. Miŝic B, Betzel RF, De Reus MA, Van Den Heuvel MP, Berman MG, McIntosh AR, et al. Network-Level Structure-Function Relationships in Human Neocortex. Cerebral Cortex. 2016;26: 3285–3296. doi:10.1093/CERCOR/BHW089

24. Suárez LE, Markello RD, Betzel RF, Misic B. Linking Structure and Function in Macroscale Brain Networks. Trends Cogn Sci. 2020;24: 302–315. doi:10.1016/J.TICS.2020.01.008

25. Vázquez-Rodríguez B, Suárez LE, Markello RD, Shafiei G, Paquola C, Hagmann P, et al. Gradients of structure–function tethering across neocortex. Proc Natl Acad Sci U S A 2019;116: 21219–21227. doi:10.1073/PNAS.1903403116/SUPPL_FILE/PNAS.1903403116.SAPP.PDF

26. Park B yong, Vos de Wael R, Paquola C, Larivière S, Benkarim O, Royer J, et al. Signal diffusion along connectome gradients and inter-hub routing differentially contribute to dynamic human brain function. Neuroimage. 2021;224: 117429. doi:10.1016/J.NEUROIMAGE.2020.117429

27. Becker CO, Pequito S, Pappas GJ, Miller MB, Grafton ST, Bassett DS, et al. Spectral mapping of brain functional connectivity from diffusion imaging. Scientific Reports 2018 8:1. 2018;8: 1–15. doi:10.1038/s41598-017-18769-x

28. Rosenthal G, Váša F, Griffa A, Hagmann P, Amico E, Goñi J, et al. Mapping higher-order relations between brain structure and function with embedded vector representations of connectomes. Nature Communications 2018 9:1. 2018;9: 1–12. doi:10.1038/s41467-018-04614-w

29. Wang P, Kong R, Kong X, Liégeois R, Orban C, Deco G, et al. Inversion of a large-scale circuit model reveals a cortical hierarchy in the dynamic resting human brain. Tropical and Subtropical Agroecosystems. 2019;21. doi:10.1126/SCIADV.AAT7854/SUPPL_FILE/AAT7854_SM.PDF

30. Deco G, Ponce-Alvarez A, Mantini D, Romani GL, Hagmann P, Corbetta M. Resting-State Functional Connectivity Emerges from Structurally and Dynamically Shaped Slow Linear Fluctuations. Journal of Neuroscience. 2013;33: 11239–11252. doi:10.1523/JNEUROSCI.1091-13.2013

31. Breakspear M. Dynamic models of large-scale brain activity. Nature Neuroscience 2017 20:3. 2017;20: 340–352. doi:10.1038/nn.4497

32. Miŝic B, Betzel RF, De Reus MA, Van Den Heuvel MP, Berman MG, McIntosh AR, et al. Network-Level Structure-Function Relationships in Human Neocortex. Cereb Cortex. 2016;26: 3285–3296. doi:10.1093/CERCOR/BHW089

33. Honey CJ, Sporns O, Cammoun L, Gigandet X, Thiran JP, Meuli R, et al. Predicting human resting-state functional connectivity from structural connectivity. Proc Natl Acad Sci U S A. 2009;106: 2035–2040. doi:10.1073/PNAS.0811168106/SUPPL_FILE/0811168106SI.PDF

34. Goni J, Van Den Heuvel MP, Avena-Koenigsberger A, De Mendizabal NV, Betzel RF, Griffa A, et al. Resting-brain functional connectivity predicted by analytic measures of network communication. Proc Natl Acad Sci U S A. 2014;111: 833–838. doi:10.1073/PNAS.1315529111/SUPPL_FILE/PNAS.201315529SI.PDF

35. Damoiseaux JS, Greicius MD. Greater than the sum of its parts: a review of studies combining structural connectivity and resting-state functional connectivity. Brain Struct Funct. 2009;213: 525–533. doi:10.1007/S00429-009-0208-6

36. Seguin C, Razi A, Zalesky A. Inferring neural signalling directionality from undirected structural connectomes. Nature Communications 2019 10:1. 2019;10: 1–13. doi:10.1038/s41467-019-12201-w

37. Suárez LE, Markello RD, Betzel RF, Misic B. Linking Structure and Function in Macroscale Brain Networks. Trends Cogn Sci. 2020;24: 302–315. doi:10.1016/J.TICS.2020.01.008

38. Benkarim O, Paquola C, Park B, Royer J, Rodríguez-Cruces R, Vos de Wael R, et al. A Riemannian approach to predicting brain function from the structural connectome. Neuroimage. 2022;257: 119299. doi:https://doi.org/10.1016/j.neuroimage.2022.119299

39. Park BY, Bethlehem RAI, Paquola C, Larivière S, Rodríguez-Cruces R, Vos de Wael R, et al. An expanding manifold in transmodal regions characterizes adolescent reconfiguration of structural connectome organization. Elife. 2021;10. doi:10.7554/ELIFE.64694

40. Duperron M-G, Knol MJ, Le Grand Q, Evans TE, Mishra A, Tsuchida A, et al. Genomics of perivascular space burden unravels early mechanisms of cerebral small vessel disease. Nature Medicine 2023 29:4. 2023;29: 950–962. doi:10.1038/s41591-023-02268-w

41. Park B yong, Kebets V, Larivière S, Hettwer MD, Paquola C, van Rooij D, et al. Multiscale neural gradients reflect transdiagnostic effects of major psychiatric conditions on cortical morphology. Communications Biology 2022 5:1. 2022;5: 1–14. doi:10.1038/s42003-022-03963-z

42. Bo T, Li J, Hu G, Zhang G, Wang W, Lv Q, et al. Brain-wide and cell-specific transcriptomic insights into MRI-derived cortical morphology in macaque monkeys. Nature Communications 2023 14:1. 2023;14: 1–15. doi:10.1038/s41467-023-37246-w

43. Park BY, Paquola C, Bethlehem RAI, Benkarim O, Mišić B, Smallwood J, et al. Adolescent development of multiscale structural wiring and functional interactions in the human connectome. Proc Natl Acad Sci U S A. 2022;119: e2116673119. doi:10.1073/PNAS.2116673119/SUPPL_FILE/PNAS.2116673119.SD01.DOCX

44. Paquola C, Bethlehem RA, Seidlitz J, Wagstyl K, Romero-Garcia R, Whitaker KJ, et al. Shifts in myeloarchitecture characterise adolescent development of cortical gradients. Elife. 2019;8. doi:10.7554/ELIFE.50482

45. Park B yong, Hong SJ, Valk SL, Paquola C, Benkarim O, Bethlehem RAI, et al. Differences in subcortico-cortical interactions identified from connectome and microcircuit models in autism. Nature Communications 2021 12:1. 2021;12: 1–15. doi:10.1038/s41467-021-21732-0

46. Hettwer MD, Larivière S, Park BY, van den Heuvel OA, Schmaal L, Andreassen OA, et al. Coordinated cortical thickness alterations across six neurodevelopmental and psychiatric disorders. Nature Communications 2022 13:1. 2022;13: 1–14. doi:10.1038/s41467-022-34367-6

47. Van Essen DC, Smith SM, Barch DM, Behrens TEJ, Yacoub E, Ugurbil K. The WU-Minn Human Connectome Project: An overview. Neuroimage. 2013;80: 62–79. doi:https://doi.org/10.1016/j.neuroimage.2013.05.041

48. Thomas Yeo BT, Krienen FM, Sepulcre J, Sabuncu MR, Lashkari D, Hollinshead M, et al. The organization of the human cerebral cortex estimated by intrinsic functional connectivity. J Neurophysiol. 2011;106: 1125. doi:10.1152/JN.00338.2011

49. Desikan RS, Ségonne F, Fischl B, Quinn BT, Dickerson BC, Blacker D, et al. An automated labeling system for subdividing the human cerebral cortex on MRI scans into gyral based regions of interest. Neuroimage. 2006;31: 968–980. doi:10.1016/j.neuroimage.2006.01.021

50. Vos de Wael R, Benkarim O, Paquola C, Lariviere S, Royer J, Tavakol S, et al. BrainSpace: a toolbox for the analysis of macroscale gradients in neuroimaging and connectomics datasets. Communications Biology 2020 3:1. 2020;3: 1–10. doi:10.1038/s42003-020-0794-7

51. Margulies DS, Ghosh SS, Goulas A, Falkiewicz M, Huntenburg JM, Langs G, et al. Situating the default-mode network along a principal gradient of macroscale cortical organization. Proc Natl Acad Sci U S A. 2016;113: 12574–12579. doi:10.1073/PNAS.1608282113/SUPPL_FILE/PNAS.201608282SI.PDF

52. Larivière S, Bayrak Ş, Vos de Wael R, Benkarim O, Herholz P, Rodriguez-Cruces R, et al. BrainStat: A toolbox for brain-wide statistics and multimodal feature associations. Neuroimage. 2023;266: 119807. doi:10.1016/J.NEUROIMAGE.2022.119807

53. Amunts K, Lepage C, Borgeat L, Mohlberg H, Dickscheid T, Rousseau MÉ, et al. BigBrain: an ultrahigh-resolution 3D human brain model. Science. 2013;340: 1472–1475. doi:10.1126/SCIENCE.1235381

54. ArnatkevicLiūtė A, Fulcher BD, Fornito A. A practical guide to linking brain-wide gene expression and neuroimaging data. Neuroimage. 2019;189: 353–367. doi:10.1016/J.NEUROIMAGE.2019.01.011

55. Markello RD, Arnatkevičiūtė A, Poline JB, Fulcher BD, Fornito A, Misic B. Standardizing workflows in imaging transcriptomics with the Abagen toolbox. Elife. 2021;10. doi:10.7554/ELIFE.72129

56. Hattox AM, Nelson SB. Layer V neurons in mouse cortex projecting to different targets have distinct physiological properties. J Neurophysiol. 2007;98: 3330–3340. doi:10.1152/JN.00397.2007

57. Larkum M. A cellular mechanism for cortical associations: an organizing principle for the cerebral cortex. Trends Neurosci. 2013;36: 141–151. doi:10.1016/J.TINS.2012.11.006

58. Anderson SA, Classey JD, Condé F, Lund JS, Lewis DA. Synchronous development of pyramidal neuron dendritic spines and parvalbumin-immunoreactive chandelier neuron axon terminals in layer III of monkey prefrontal cortex. Neuroscience. 1995;67: 7–22. doi:10.1016/0306-4522(95)00051-J

59. Kim EJ, Juavinett AL, Kyubwa EM, Jacobs MW, Callaway EM. Three Types of Cortical Layer 5 Neurons That Differ in Brain-wide Connectivity and Function. Neuron. 2015;88: 1253–1267. doi:10.1016/j.neuron.2015.11.002

60. Briggs F. Organizing principles of cortical layer 6. Front Neural Circuits. 2010;4: 3. doi:10.3389/NEURO.04.003.2010/BIBTEX

61. Valk SL, Xu T, Margulies DS, Masouleh SK, Paquola C, Goulas A, et al. Shaping brain structure: Genetic and phylogenetic axes of macroscale organization of cortical thickness. Sci Adv. 2020;6. doi:10.1126/SCIADV.ABB3417/SUPPL_FILE/ABB3417_SM.PDF

62. Valk SL, Xu T, Paquola C, Park B yong, Bethlehem RAI, Vos de Wael R, et al. Genetic and phylogenetic uncoupling of structure and function in human transmodal cortex. Nature Communications 2022 13:1. 2022;13: 1–17. doi:10.1038/s41467-022-29886-1

63. Buckner RL, Krienen FM. The evolution of distributed association networks in the human brain. Trends Cogn Sci. 2013;17: 648–665. doi:10.1016/J.TICS.2013.09.017

64. Hill J, Inder T, Neil J, Dierker D, Harwell J, Van Essen D. Similar patterns of cortical expansion during human development and evolution. Proc Natl Acad Sci U S A. 2010;107: 13135–13140. doi:10.1073/PNAS.1001229107/SUPPL_FILE/PNAS.201001229SI.PDF

65. Ottino-González J, Baggio HC, Jurado MÁ, Segura B, Caldú X, Prats-Soteras X, et al. Alterations in Brain Network Organization in Adults with Obesity as Compared with Healthy-Weight Individuals and Seniors. Psychosom Med. 2021;83: 700–706. doi:10.1097/PSY.0000000000000952

66. Gómez-Apo E, García-Sierra A, Silva-Pereyra J, Soto-Abraham V, Mondragón-Maya A, Velasco-Vales V, et al. A Postmortem Study of Frontal and Temporal Gyri Thickness and Cell Number in Human Obesity. Obesity. 2018;26: 94–102. doi:https://doi.org/10.1002/oby.22036

67. Burger KS, Stice E. Greater striatopallidal adaptive coding during cue–reward learning and food reward habituation predict future weight gain. Neuroimage. 2014;99: 122–128. doi:https://doi.org/10.1016/j.neuroimage.2014.05.066

68. Rodríguez-Rodríguez R, Miralpeix C. Hypothalamic Regulation of Obesity. International Journal of Molecular Sciences 2021, Vol 22, Page 13459. 2021;22: 13459. doi:10.3390/IJMS222413459

69. Jennings JH, Rizzi G, Stamatakis AM, Ung RL, Stuber GD. The Inhibitory Circuit Architecture of the Lateral Hypothalamus Orchestrates Feeding. Science (1979). 2013;341: 1517–1521. doi:10.1126/science.1241812

70. Clarke RE, Verdejo-Garcia A, Andrews ZB. The role of corticostriatal–hypothalamic neural circuits in feeding behaviour: implications for obesity. J Neurochem. 2018;147: 715–729. doi:https://doi.org/10.1111/jnc.14455

71. Jenkinson M, Beckmann CF, Behrens TEJ, Woolrich MW, Smith SM. FSL. Neuroimage. 2012;62: 782–790. doi:10.1016/J.NEUROIMAGE.2011.09.015

72. Fischl B. FreeSurfer. Neuroimage. 2012;62: 774–781. doi:10.1016/J.NEUROIMAGE.2012.01.021

73. Glasser MF, Sotiropoulos SN, Wilson JA, Coalson TS, Fischl B, Andersson JL, et al. The minimal preprocessing pipelines for the Human Connectome Project. Neuroimage. 2013;80: 105–124. doi:https://doi.org/10.1016/j.neuroimage.2013.04.127

74. Dale AM, Fischl B, Sereno MI. Cortical Surface-Based Analysis: I. Segmentation and Surface Reconstruction. Neuroimage. 1999;9: 179–194. doi:10.1006/NIMG.1998.0395

75. Fischl B, Sereno MI, Dale AM. Cortical Surface-Based Analysis: II: Inflation, Flattening, and a Surface-Based Coordinate System. Neuroimage. 1999;9: 195–207. doi:10.1006/NIMG.1998.0396

76. Fischl B, Sereno MI, Tootell RBH, Dale AM. High-Resolution Intersubject Averaging and a Coordinate System for the Cortical Surface. Hum Brain Mapping. 1999;8: 272–284. doi:10.1002/(SICI)1097-0193(1999)8:4

77. Van Essen DC, Glasser MF, Dierker DL, Harwell J, Coalson T. Parcellations and Hemispheric Asymmetries of Human Cerebral Cortex Analyzed on Surface-Based Atlases. Cerebral Cortex. 2012;22: 2241–2262. doi:10.1093/CERCOR/BHR291

78. Glasser MF, Coalson TS, Robinson EC, Hacker CD, Harwell J, Yacoub E, et al. A multi-modal parcellation of human cerebral cortex. Nature 2016 536:7615. 2016;536: 171–178. doi:10.1038/nature18933

79. Salimi-Khorshidi G, Douaud G, Beckmann CF, Glasser MF, Griffanti L, Smith SM. Automatic denoising of functional MRI data: Combining independent component analysis and hierarchical fusion of classifiers. Neuroimage. 2014;90: 449–468. doi:10.1016/J.NEUROIMAGE.2013.11.046

80. Tournier JD, Smith R, Raffelt D, Tabbara R, Dhollander T, Pietsch M, et al. MRtrix3: A fast, flexible and open software framework for medical image processing and visualisation. Neuroimage. 2019;202: 116137. doi:10.1016/J.NEUROIMAGE.2019.116137

81. Smith RE, Tournier JD, Calamante F, Connelly A. Anatomically-constrained tractography: Improved diffusion MRI streamlines tractography through effective use of anatomical information. Neuroimage. 2012;62: 1924–1938. doi:10.1016/J.NEUROIMAGE.2012.06.005

82. Jeurissen B, Tournier JD, Dhollander T, Connelly A, Sijbers J. Multi-tissue constrained spherical deconvolution for improved analysis of multi-shell diffusion MRI data. Neuroimage. 2014;103: 411–426. doi:10.1016/J.NEUROIMAGE.2014.07.061

83. Tournier J, Calamante F, Connelly A. Improved probabilistic streamlines tractography by 2 nd order integration over fibre orientation distributions. 2009.

84. Smith RE, Tournier JD, Calamante F, Connelly A. SIFT2: Enabling dense quantitative assessment of brain white matter connectivity using streamlines tractography. Neuroimage. 2015;119: 338–351. doi:10.1016/J.NEUROIMAGE.2015.06.092

85. Schaefer A, Kong R, Gordon EM, Laumann TO, Zuo X-N, Holmes AJ, et al. Local-Global Parcellation of the Human Cerebral Cortex from Intrinsic Functional Connectivity MRI. Cerebral Cortex. 2018;28: 3095–3114. doi:10.1093/cercor/bhx179

86. Thompson WH, Fransson P. On Stabilizing the Variance of Dynamic Functional Brain Connectivity Time Series. Brain Connect. 2016;6: 735–746. doi:10.1089/BRAIN.2016.0454/ASSET/IMAGES/LARGE/FIGURE6.JPEG

87. Coifman RR, Lafon S. Diffusion maps. Appl Comput Harmon Anal. 2006;21: 5–30. doi:10.1016/J.ACHA.2006.04.006

88. Langs G, Golland P, Ghosh SS. Predicting activation across individuals with resting-state functional connectivity based multi-atlas label fusion. Lecture Notes in Computer Science (including subseries Lecture Notes in Artificial Intelligence and Lecture Notes in Bioinformatics). 2015;9350: 313–320. doi:10.1007/978-3-319-24571-3_38/COVER

89. Benjamini Y, Hochberg Y. Controlling the False Discovery Rate: A Practical and Powerful Approach to Multiple Testing. Journal of the Royal Statistical Society: Series B (Methodological). 1995;57: 289–300. doi:10.1111/J.2517-6161.1995.TB02031.X

90. Paquola C, Vos De Wael R, Wagstyl K, Bethlehem RAI, Hong SJ, Seidlitz J, et al. Microstructural and functional gradients are increasingly dissociated in transmodal cortices. PLoS Biol. 2019;17: e3000284. doi:10.1371/JOURNAL.PBIO.3000284

91. Alexander-Bloch AF, Shou H, Liu S, Satterthwaite TD, Glahn DC, Shinohara RT, et al. On testing for spatial correspondence between maps of human brain structure and function. Neuroimage. 2018;178: 540–551. doi:10.1016/J.NEUROIMAGE.2018.05.070

92. Hawrylycz MJ, Lein ES, Guillozet-Bongaarts AL, Shen EH, Ng L, Miller JA, et al. An anatomically comprehensive atlas of the adult human brain transcriptome. Nature 2012 489:7416. 2012;489: 391–399. doi:10.1038/nature11405

93. Dougherty JD, Schmidt EF, Nakajima M, Heintz N. Analytical approaches to RNA profiling data for the identification of genes enriched in specific cells. Nucleic Acids Res. 2010;38: 4218–4230. doi:10.1093/NAR/GKQ130

